# Enhanced Stability of Non-Canonical NPC2 in the symbiosome supports coral-algal symbiosis

**DOI:** 10.1101/399766

**Authors:** Elizabeth A. Hambleton, Victor A.S. Jones, Ira Maegele, David K vaskoff, Timo Sachsenheimer, Annika Guse

## Abstract

Cnidarians such as reef-building corals depend upon nutrient transfer from intracellular symbionts, but the mechanisms and evolution of this process remain unknown. Homologues of the conserved cholesterol binder Niemann-Pick Type C2 (NPC2) in cnidarians are implicated in the transfer of sterol from symbionts. Here, we show that symbionts transfer bulk sterols to the host, host sterol utilization is plastic, and pharmacological inhibition of sterol trafficking disrupts symbiosis. Having undergone an anthozoan-specific expansion, “non-canonical” NPC2s respond to symbiosis and accumulate over time at the lysosomal-like organelle in which the symbiont resides (“symbiosome”). We demonstrate that both a non- and canonical *Aiptasia* NPC2 bind symbiont-produced sterols, yet only the non-canonical homologue exhibits increased stability at low pH. We propose that symbiotic cnidarians adapted pre-existing cholesterol-trafficking machinery to function in the highly acidic symbiosome environment, allowing corals to dominate nutrient-poor shallow tropical seas worldwide.

---

Many plants and animals cultivate symbioses with microorganisms for nutrient exchange. Cnidarians, such as reef-building corals and anemones, establish an ecologically critical endosymbiosis with photosynthetic dinoflagellate algae (*Symbiodinium* spp.) [*Douglas*]. Their symbionts reside within endo/lysosomal-like organelles termed symbiosomes and transfer photosynthetic products to their hosts [*Muscatine, Yellowlees*]. In addition to sugars, lipids are among the key products transferred [*Crossland, Battey, Revel*]. Cnidarians are sterol auxotrophs [*Baumgarten, Gold*] that must acquire these essential compounds from diet and/or symbionts [*Goad*]. Dinoflagellates synthesize various sterols, many of which are found in symbiotic cnidarians [*Withers, Bohlin, Ciereszko*], yet how sterol transfer occurs is undetermined.

The highly conserved Niemann-Pick Type C2 protein (NPC2) is a small, soluble lysosomal protein that facilitates cholesterol egress to the cytoplasm [*Vance*]. Symbiotic cnidarians contain an additional subset of NPC2 homologues that form a separate group based on sequence, designated “non-canonical” NPC2s [*Dani14, Lehnert*]. Non-canonical NPC2s are consistently up-regulated in symbiosis in *Aiptasia* and *Anemonia* anemones [*Kuo, Ganot, Dani14, Lehnert, Wolfowicz*] and these symbiosis-responding NPC2s localize around the symbiosome [*Dani14, Dani17*]. Therefore, the current hypothesis in the field is that non-canonical NPC2s specifically facilitate transfer of symbiont-produced sterols in cnidarian-algal symbiosis [*Baumgarten, Dani17, Revel*, *Wolfowicz*]. However, NPC2s may serve other purposes, e.g. signalling [*Baumgarten, Dani17*], and mechanistic analyses of NPC2 function are lacking. We therefore sought to determine the physiological roles of cnidarian NPC2s, which might explain their persistence throughout evolution.

We first determined the full genomic complement of NPC2 homologues in symbiotic cnidarians and specific metazoans, uncovering several previously unidentified homologues in the reef-building corals and other taxa (asterisks, Fig. 1A). We conducted a Bayesian reconstruction of phylogeny, recovering the division between canonical and non-canonical NPC2s and corresponding stereotypic intron/exon structures (Fig. 1A). We found that non-canonical NPC2 homologues are thus far confined to cnidarians within the anthozoan class, as they did not appear in the earlier-branching sponge *Amphimedon* nor in the hydrozoans *Hydra magnipapillata* and *Hydractinia echinata*. Notably, the degree of NPC2 gene expansion appeared to correlate with symbiotic state: the symbiotic anthozoans (*Aiptasia*, *Acropora, Montastrea*) have several non-canonical NPC2 homologues (3, 3, and 2, respectively). In contrast, the non-symbiotic anemone *Nematostella* displays evolutionary traces of a single non-canonical NPC2, which either failed to expand or underwent higher loss (Fig. 1A).

**Figure 1.**
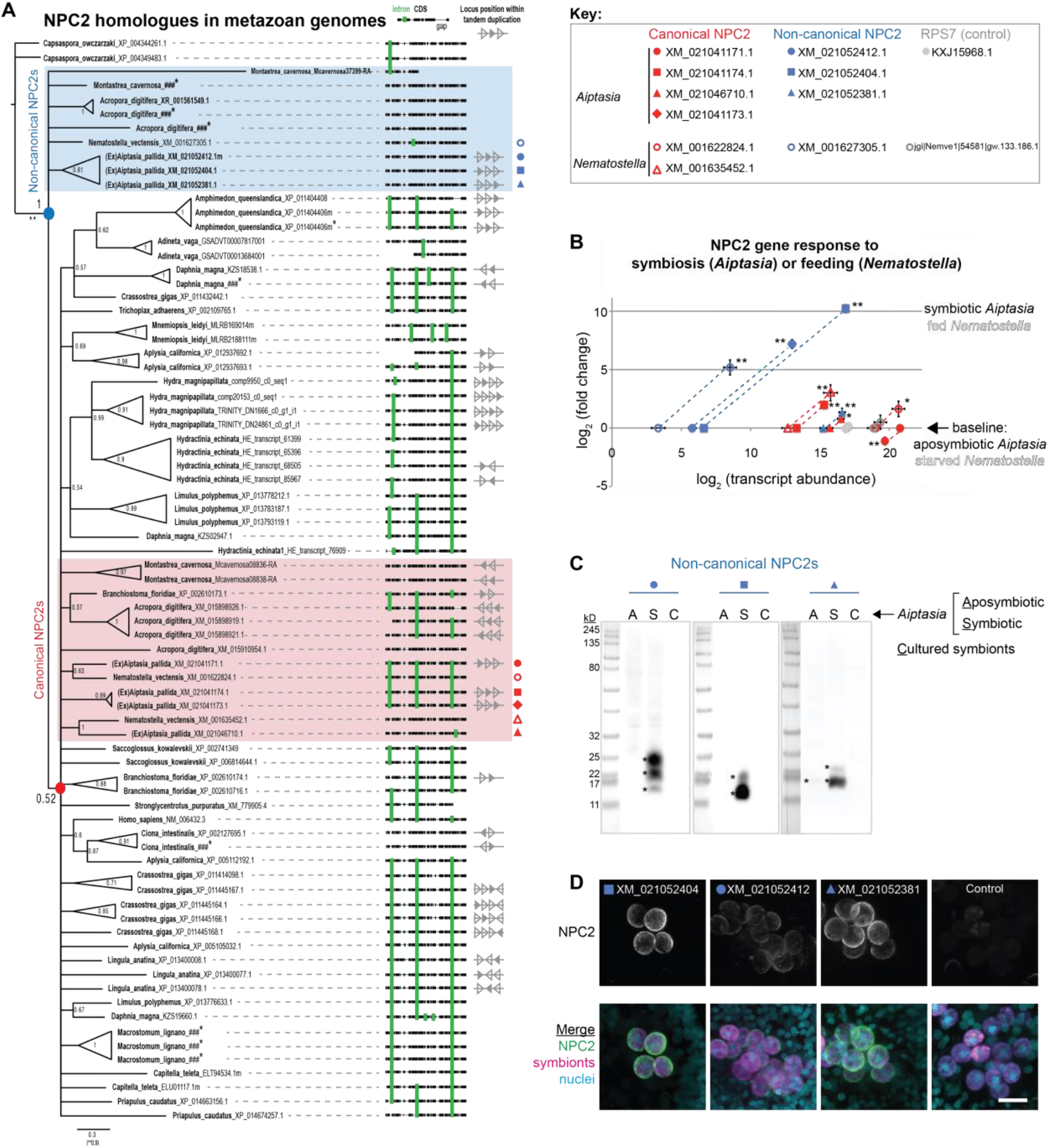
Symbiotic anthozoans have extensive expansion of NPC2s that respond to symbiosis. **A.** Consensus phylogeny from Bayesian analysis (MrBayes) of NPC2 coding sequences in metazoan genomes with anthozoan non-canonical (blue shading) and canonical (red shading) homologues. Also shown are alignments with intron/exon boundaries (green bars) and tandem duplication of NPC2 loci (where genome assemblies allow). Node values, posterior probabilities. Asterisks, new homologues from this study. Source data, see Methods. **B.** Gene expression by RT-qPCR of canonical (red symbols) and non-canonical (blue symbols) NPC2s and 40S ribosomal subunit (RPS7, grey symbols). Filled symbols: Log_2_ (fold change) in symbiotic versus aposymbiotic (on x-axis) *Aiptasia*. Average of 6 samples per condition, 6 animals per sample, each in technical duplicate. Open symbols: Log_2_ (fold change) in fed versus starved (on x-axis) *Nematostella*. Average of 2 animals per sample, 2 samples per condition, technical duplicates. For all, average value ±SD (error bars). Statistical comparisons per gene of symbiotic (/fed) to aposymbiotic (/starved) by Bayesian modeling, *P<0.05, **P<0.005 (RPS7 not significant). **C.** Homogenates of *Aiptasia* symbiotic and aposymbiotic adults detected with affinity-purified antibodies to non-canonical *Aiptasia* NPC2 homologues. Asterisks, NPC2 glycoforms (see Fig. 3A). **D.** Immunofluorescence (IF) of non-canonical NPC2s in *Aiptasia* larvae containing intracellular symbionts of *Symbiodinium* strain SSB01 [*Xiang*]. Larvae are 14 d post-fertilization (dpf) and 11 d post-infection (dpi). Merge channels: NPC2, secondary antibody Alexa488-anti-rabbit IgG; Nuclei, Hoechst; Symbionts, red autofluorescence of photosynthetic machinery. Control, secondary antibody only. Scale bar, 10 μm.

To investigate the function of NPC2s in symbiosis, we analysed their expression in the tropical sea anemone *Aiptasia*, a powerful emerging model of cnidarian-algal symbiosis [*Neubauer, Tolleter*]. *Aiptasia* contains three non-canonical NPC2 homologues. Two of these non-canonical NPC2 homologues displayed substantially high expression at the transcript and protein levels in symbiotic animals but not aposymbiotic animals (closed blue symbols, Fig. 1B and 1C). The third non-canonical NPC2 homologue was highly expressed in both symbiotic and aposymbiotic animals (Fig. 1B and 1C). Conversely, canonical NPC2s were highly expressed in both symbiotic and aposymbiotic animals (closed red symbols, Fig. 1B). Likewise, the non-symbiotic anemone *Nematostella* exhibited ubiquitously high expression of canonical NPC2 genes (open red symbols, Fig. 1B), whereas the non-canonical NPC2 gene was highly expressed only upon feeding (open blue symbols, Fig. 1B). Aposymbiotic embryos of the symbiotic coral *Acropora*, as well as *Nematostella* embryos, contained maternally provided canonical NPC2 transcripts, suggesting that these are required for development (Fig. S1). The non-canonical NPC2s decorated intracellular symbionts *in vivo* in *Aiptasia* (Fig. 1D), consistent with previous data in *Anemonia viridis [Dani14, Dani17*]. Together, these data suggest evolutionary conserved roles for canonical and non-canonical NPC2, with a continuous requirement for high levels of canonical NPC2s, but a nutrient-dependent requirement for non-canonical NPC2s in the lysosomal-like symbiosome.

Notably, several canonical NPC2s in *Aiptasia* (XM_021046710, XM_021041174) and *Nematostella* (XM_0016335452) may be ‘in transition’ to becoming non-canonical: they were expressed at intermediate abundances between the two groups, and they responded to symbiosis (*Aiptasia*) or feeding (*Nematostella*) (red square and triangles, Fig. 1B and Fig. S1). Further, some of their intron/exon structures reflected those of the non-canonical group (red triangles, Fig. 1A). This suggests that a continuous diversification of NPC2 variants may be advantageous for these species.

Sterol compositions of the *Symbiodinium* symbionts are complex and vary by strain [*Withers, Bohlin, Ciereszko*]. To test whether changes in symbiont-provided nutrients affect NPC2 gene expression, we used gas chromatography/mass spectrometry (GC/MS) to semi-quantitatively profile relative sterol abundances in compatible symbiont strains [*Xiang, Hambleton*] and related this to NPC2 gene expression in various host lines. We found that the sterol profile is a direct consequence of the symbiont in a defined host genotype (Fig. 2A). For example, *Aiptasia* CC7 hosting *Symbiodinium* strain SSA01 contained a large proportion of stigmasterol-like sterol. In contrast, the same animal line hosting strain SSB01 contained minimal stigmasterol-like derivatives yet a sizable proportion of the unique sterol gorgosterol (Fig. 2A), characterized by an unusual cyclopropyl group [*Ciereszko*] (Fig. S2). Further, the genetically distinct *Aiptasia* lines H2 and CC7 [*Grawunder*] showed almost identical sterol profiles when hosting the same symbiont strain (Fig. 2A). Importantly, *Aiptasia* lines showed similar patterns of NPC2 gene expression in each of these combinations relative to their aposymbiotic counterparts (Fig. 2B). These data suggest that host sterol uptake is independent of the particular sterol suite provided, and that non-canonical NPC2 genes respond robustly and consistently to different symbionts.

**Figure 2.**
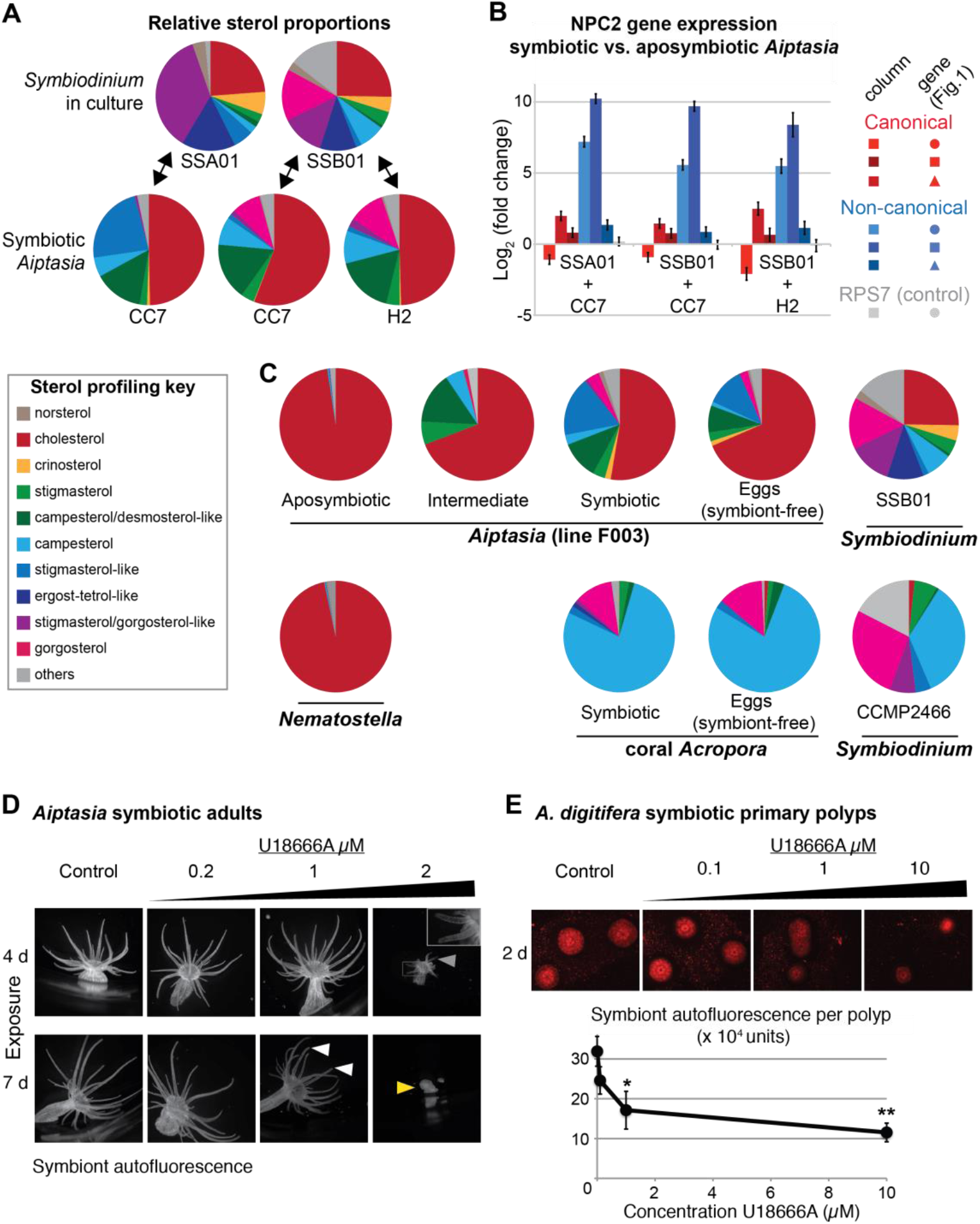
NPC2s respond consistently to symbionts, and symbiont-produced sterol acceptance is plastic in the host yet critical for tissue homeostasis and symbiosis. **A.** Gas chromatography/mass spectrometry (GC/MS)-generated sterol profiles of *Symbiodinium* strains [*Xiang*] in culture (upper row) and in symbiosis with adults in different *Aiptasia* host strains [*Grawunder*] (lower row). Relative composition (%) of each sterol in key, below. Shown is representative of n=3 (*Aiptasia, Symbiodinium*) or n=2 (*Acropora, Nematostella*) samples. **B.** NPC2 gene expression by qPCR in the various *Aiptasia*/*Symbiodinium* host/symbiont combinations in ***A***. Gene symbols as in Fig. 1. **C.** GC/MS-generated sterol profiles of the given organisms, details as in ***A***. Apo-and non-symbiotic animals were fed *Artemia* brine shrimp comprising nearly only cholesterol [*Tolosa*]. “Intermediate” were symbiotic *Aiptasia* more recently cured of symbionts. *Aiptasia* strain F003 hosts *Symbiodinium* strains SSA01 and SSB01 [*Xiang*]. *Acropora digitifera* endogenous *Symbiodinium* are currently uncultured but are closely related to the cultured strain CCMP2466 (see Methods). **D.** Fluorescence micrographs of adult polyps of *Aiptasia* strain F003 exposed to U18666A or DMSO negative control (vol. equiv. to 10µM addition). Autofluorescence is symbiont photosynthetic machinery. Representative images shown of n=6 polyps per condition, from one of two replicate experiments. Arrows: grey, symbiont loss in tentacles; white, early indications of tissue disruption (shortened tentacles); yellow, polyp with reduced tentacles and body mass. **E.** Fluorescence micrographs of *Acropora digitifera* juvenile primary polyps hosting *Symbiodinium* strain SSB01 exposed to U18666A or negative control, as in ***D***. Representative images shown of n=5 polyps across duplicate wells (n=4 for 10µM). Quantification of total fluorescence per polyp minus adjacent background. Average values ±SEM (error bars). Statistical comparisons to DMSO negative control (Student’s *t*-test: *P<0.05, **P<0.005).

Indeed, we found that cnidarians display remarkable flexibility in sterol composition depending on nutrient input: aposymbiotic *Aiptasia* and their non-symbiotic relatives *Nematostella* contained almost exclusively cholesterol, reflecting their *Artemia*-based diet (Fig. 2C). However, aposymbiotic animals more recently cured of their symbionts (“intermediate”) contain traces of the former symbiont input together with dietary cholesterol (Fig. 2C). Crucially, naturally aposymbiotic eggs of *Aiptasia* exhibited sterol patterns very similar to their symbiotic parents, confirming that these symbiont-produced sterols are indeed transferred to host tissues (Fig. 2C). Likewise, the sterol profiles of the coral *Acropora* and their non-symbiotic eggs reflected their symbionts. Strikingly, field-collected corals were nearly devoid of cholesterol (Fig. 2C), suggesting that when food is scarce, symbiotic cnidarians are capable of functionally substituting cholesterol with symbiont-produced sterols. Thus, the sterol composition of symbiotic cnidarians is highly plastic and adaptable to environmental conditions, including food availability.

In accordance with the functionally important role of symbiont-produced sterols in the host, we found that disruption of sterol trafficking in symbiosis has profound effects. To evaluate the role of sterol trafficking in symbiosis, we utilized the drug U18666A, which blocks sterol egress from the lysosome [*Liscum, Cenedella*]. U18666A is a competitive inhibitor of NPC1, a binding partner of NPC2 that is required for efficient cholesterol egress [*Lu*]. Symbiotic adult *Aiptasia* showed a dose-and duration-dependent disruption of host physiology and symbiosis in response to U18666A (Fig. 2D), including loss of anemone tissue and shortening of tentacles, as well as loss of symbionts in the tissue (inset, Fig. 2D). Additionally, we exposed *A. digitifera* juvenile primary polyps stably hosting the same symbionts (*Symbiodinium* strain SSB01) to increasing concentrations of U18666A (Fig. 2E), which to our knowledge represents the first reported functional disruption of sterol transport in coral symbiosis. We quantified symbiont loads, and again observed a U18666A dose-dependent loss of symbionts in the polyps (Fig. 2E), mirroring the effects in *Aiptasia*. Thus, inhibition of sterol transport affects symbiosis stability, leading to loss of symbionts (“bleaching”). Further, the disruption of sterol transport compromises host tissues, emphasizing the importance of sterols in tissue homeostasis.

Mammalian NPC2s function in lysosomal sterol binding, and our evidence so far suggests that non-canonical NPC2s play a specific role in the symbiosome. However, whether non-canonical and canonical NPC2s functionally differ – and indeed, whether cnidarian NPC2s bind any sterols – are not known. To elucidate NPC2 functions, we first turned to the structure-function characterizations of mammalian NPC2s. Canonical NPC2 has a single hydrophobic pocket that binds the side-chain of cholesterol [*Friedland*], and important residues for binding and transfer have been identified [*Ko, Xu, Wang, McCauliff*] (Fig. 3A). Using protein-wide adaptive evolution analysis of cnidarian NPC2s, we calculated per-residue evolutionary rates (Fig. S3) and we identified residues apparently under positive (diversifying) or negative (purifying) selection (Fig. 3A). We found that, overall, the hydrophobic pocket is more highly conserved in canonical NPC2 proteins (17 of 35 residues under negative selection) than in non-canonical NPC2 proteins (3 of 35 residues) in *Aiptasia*. Moreover, 5 of 12 residues under positive selection in the non-canonical NPC2s fall in this region. We therefore hypothesized that the ligand-binding pocket has diverged for a symbiosis-specific function, and therefore that non-canonical NPC2 may have evolved to bind symbiont-derived sterols more effectively than their non-canonical counterparts.

**Figure 3.**
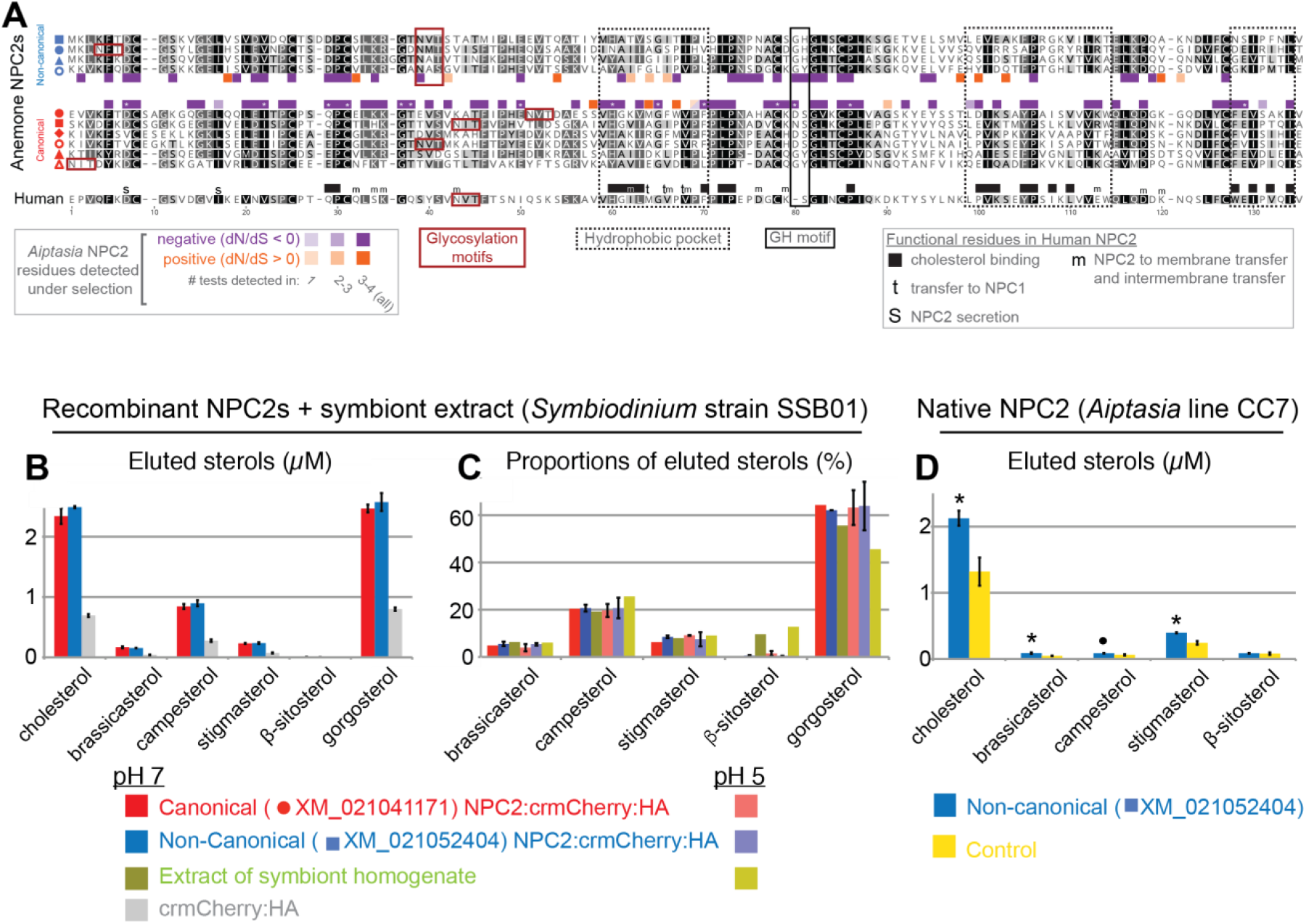
Adaptive evolution analysis of NPC2s and their binding to symbiont-produced sterols via immunoprecipitation-lipidomics. **A**. Alignment of anemone and human NPC2 proteins, with black-white shading by conservation. Highlighted are residues under positive (orange) or negative (purple) selection per NPC2 group as found in multiple tests of non-synonymous/synonymous substitution rates (dN/dS) in HyPhy [*Kosakovsky Pond_1*] (see Methods); asterisks, significant in all tests. Indicated are also several functional regions in human NPC2 [*Ko, Xu, Wang, McCauliff*]. **B.** Quantification of bound lipids in the eluates following IP of recombinant canonical and non-canonical NPC2:crmCherry:HA and negative control crmCherry:HA at pH 7 (Fig. S4). Sterols comprising <1.5% were omitted for clarity. crmCherry, lysosome-stable <u>c</u>leavage-resistant mCherry [*Huang*]. Average values ±SD (error bars) of duplicates, representative experiment of three replicate experiments shown. Statistical comparisons of each NPC2 to crmCherry negative control, all differences significant with Student’s *t*-test at P<0.01 except ß-sitosterol values. **C.** Relative proportions of NPC2-bound sterols and the corresponding symbiont extract at pH 5 and 7. Average values ±SD (error bars) of duplicates, representative of three replicate experiments shown. **D.** Immunoprecipitation (IP) of native non-canonical NPC2 from *Aiptasia* and quantification of eluted bound sterols. Control, identical reaction omitting antibody. Average values ±SEM (error bars) of duplicate samples, representative of two duplicate experiments shown. Statistical comparisons to control (Student’s *t*-test: *P<0.05, •P<0.09).

To test this hypothesis of differential NPC2-sterol binding in *Aiptasia*, we directly compared the most conserved canonical NPC2 to the non-canonical NPC2 that is most up-regulated upon symbiosis (XM_021041171 to XM_021052404, respectively). We used lipidomics to quantify lipids bound by immunoprecipitated native or recombinant NPC2s (Fig. S4) [after *Li*]. Recombinant proteins were expressed in HEK 293T cells, after which cell lysates were mixed with *Symbiodinium* SSB01 homogenates at either neutral conditions (pH 7) or acidic conditions reflecting the lysosome (pH 5). Under both conditions, canonical and non-canonical NPC2:mCherry fusion proteins bound symbiont-produced sterols significantly above the background levels of the control, mCherry alone (Fig. 3B). The relative proportions of bound sterols generally exhibited equilibrium levels with the corresponding symbiont homogenate. Thus, this assay did not detect any differential sterol binding between canonical vs. non-canonical NPC2s (Fig. 3C). To validate sterol binding by non-canonical NPC2 *in vivo*, we also immunoprecipitated the native non-canonical NPC2 and bound sterols directly from homogenates of symbiotic *Aiptasia*. Again, we detected symbiont-produced sterols above background levels, validating our heterologous system and indicating that these proteins bind sterols *in vivo* during symbiosis (Fig. 3D). These data indicate that, despite their evolutionary divergence, both types of *Aiptasia* NPC2s have the conserved function of binding sterols in lysosomal-like environments. Although we cannot rule out subtle differences in sterol binding dynamics between the two proteins, our results suggest that differential binding does not appear to be a distinguishing feature of non-canonical NPC2s. These findings are consistent with the observations that the sterol ligand and the residues lining the binding cavity tolerate variations [*Xu, Liou*].

With data suggesting both NPC2 types can bind symbiont-produced sterols, we were therefore left with the question: what is the functional advantage of localizing non-canonical NPC2s specifically in the symbiosome? We noted that non-canonical NPC2s decorate some but not all symbionts (Fig. 1D, *Dani14*, *Dani17*), suggesting that at any given time, symbiosomes are a dynamic group of specialized organelles. To gain further insight into the NPC2-decorated symbiosome dynamics, we measured the spatio-temporal regulation of non-canonical NPC2 by immunofluorescence in *Aiptasia* larvae establishing symbiosis (“infection”) with *Symbiodinium* strain SSB01 [*Hambleton, Bucher*]. Indeed, non-canonical NPC2 slowly decorated intracellular symbionts over time (Fig. 4A, Fig. S5). This localization ranged from weak ‘grainy’ patterns to stronger ‘halos’ around symbionts (arrows, Fig. 4B). We quantified infection rates, symbiont load of individual larvae, and non-canonical NPC2 signal intensity (Fig. 4C-D). We found that infection rates remained steady after removal of symbionts from the environment, whereas the proportion of larvae showing non-canonical NPC2 signal continued to increase to eventually include the majority of infected larvae (Fig. 4C). Concordantly, the proportion of symbionts within each larva surrounded by NPC2 signal also increased over time, as did the signal strength (Fig. 4C). Finally, infected larvae displaying any NPC2 signal generally contained a higher symbiont load than their infected, unlabeled counterparts (Fig. 4D). Thus, non-canonical NPC2 is increasingly expressed and recruited to symbionts over time, suggesting that non-canonical NPC2 function becomes important primarily once symbiosomes are “mature”.

**Figure 4.**
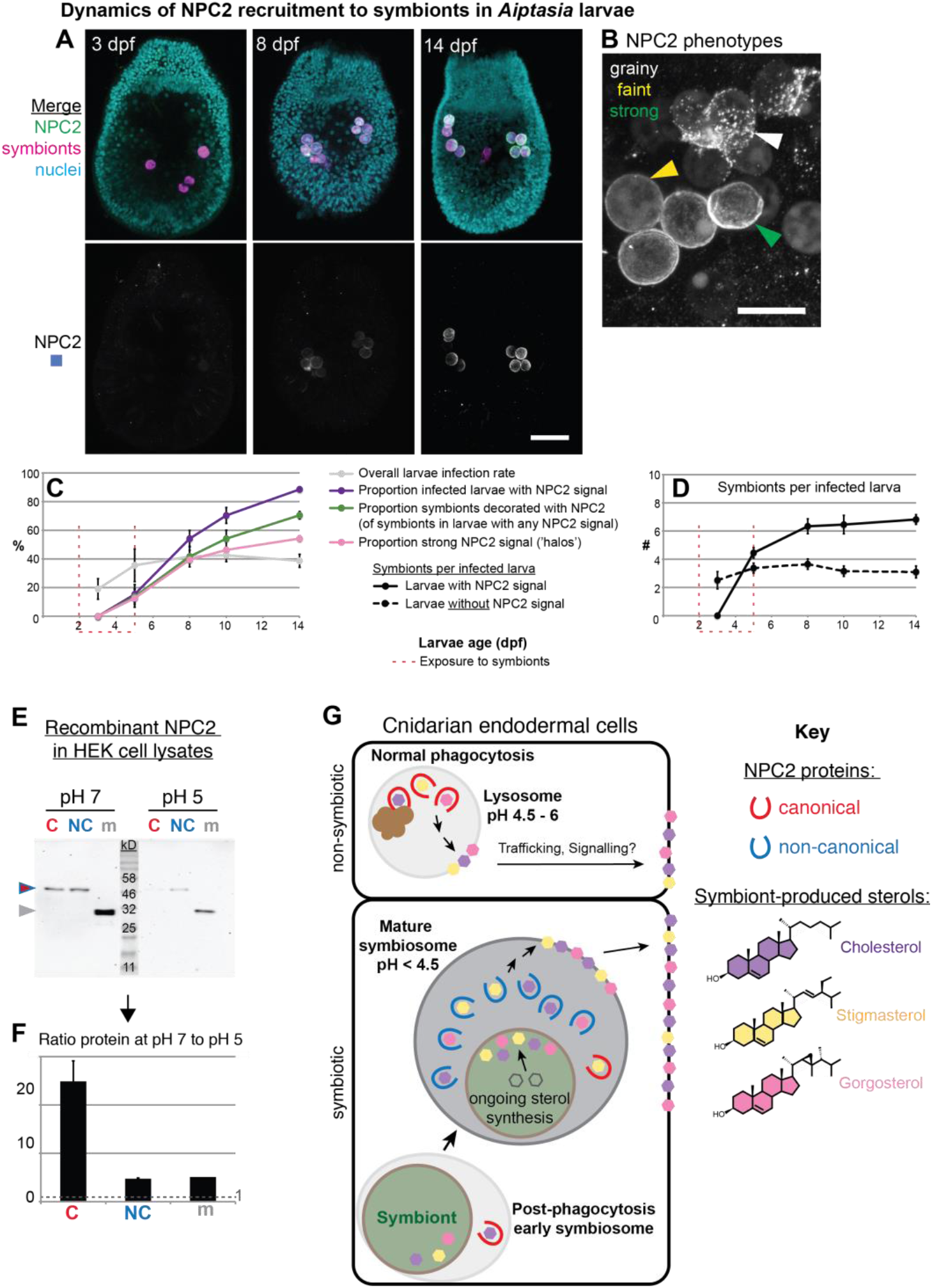
Non-canonical NPC2 is spatiotemporally regulated to mature symbiosomes and is more stable in acidic environments. **A.** Time-course of immunofluorescence of non-canonical NPC2 in *Aiptasia* larvae infected with *Symbiodinium* SSB01 from 2-5dpf. Larvae oral opening facing up. Merge channels: NPC2, secondary antibody Alexa488-anti-rabbit IgG; Nuclei, Hoechst; Symbionts, red autofluorescence of photosynthetic machinery. Control, secondary antibody only. Scale bar, 25 μm. **B.** Patterns of NPC2 labeling observed. Scale bar, 10 μm. **C-D.** Quantification of NPC2 IF time-course in ***B***. n=triplicate samples of >50 larvae per time-point. Representative of two independent experiments. Average value ±SEM (error bars). **E.** Recombinant NPC2s in the soluble fractions of HEK cell lysates. Lysate preparations were identical except for buffer pH; equivalent volumes loaded per lane. One pair of treatments in a representative experiment shown, of two duplicate experiments. **F.** Quantification of protein abundances from ***E***. Measurements taken within a Western blot, with n=2 (NPC2s) or n=1 (mCherry) measurements from different blot scans. Average values ±SEM (error bars), representative of two duplicate experiments shown. **G.** We propose a model in which symbiotic anthozoans have evolved non-canonical NPC2 homologues (blue) that are spatiotemporally regulated to specifically respond to symbiosis, including through adaptation to the extremely acidic environment of the symbiosome, the lysosomal-like organelle in which symbionts reside. In contrast, ubiquitously expressed canonical NPC2s (red) are found throughout the tissue as ‘workhorses’ in sterol trafficking. Within their respective environments, both NPC2 types bind and transport symbiont-produced sterols (purple, yellow, pink), and such trafficking is essential for membrane homeostasis of the sterol-auxotrophic hosts.

The maturing symbiosome, where non-canonical NPC2s function, remains poorly understood; however, extreme acidity is a unique characteristic of these specialized cellular compartments. Whereas lumenal pH of classic lysosomes can range from 4.7 to 6 [*Johnson*], recent work indicates that mature symbiosomes in steady-state symbiosis are even more acidic (pH ∼4) to promote efficient photosynthesis [*Barott*]. Protein activity, stability, and solubility are dependent on pH, and proteins have tailored their amino acid composition for structural and functional purposes under specific pH conditions [*Talley*]. It remains difficult to correlate pH-dependent protein properties with aspects of their sequence or structure, largely because proteins evolved under myriad physical and biological constraints [*Garcia-Moreno*]. However, we noted two features among highly conserved *Aiptasia* non-canonical NPC2 proteins: glycosylation sites (Fig. 3A) and a glycine followed by a histidine residue (Fig. 3A). Both glycosylation sites and histidines (differential protonation at low pH) can contribute to protein stability in acidic environments [*Rudd*, *Hanson*, *Culyba*].

We therefore sought to compare the stability/solubility of canonical and non-canonical NPC2 at different pH. We showed above that both types of *Aiptasia* NPC2s bound sterols at pH 7 and pH 5 (Fig. 3B-C). However, we found that at pH 5, non-canonical NPC2 was consistently more abundant in the soluble fraction than canonical NPC2 (Fig. 4E). For each type of NPC2, the ratio of soluble protein at pH 7 to that at pH 5 was always >1, indicating that both were more soluble at pH 7. However, the ratio was higher for canonical NPC2 than for non-canonical NPC2 and the control mCherry, indicating that canonical NPC2 becomes far less soluble at pH 5 (Fig. 4F). Thus, this non-canonical NPC2 appears to be more soluble/stable than the canonical NPC2 at a lower pH, characteristic of the symbiosome. It will be important in the future to experimentally test whether increased pH-stability is a common feature of non-canonical NPC2s in symbiotic cnidarians, and whether these proteins have evolved similar or different strategies to achieve this goal. The increased sequence diversity among anthozoan non-canonical NPC2 (Fig. 1A) points towards the latter.

Taken together, our data support a model in which symbiotic anthozoans have evolved symbiosis-specific NPC2 homologues that have adapted their solubility to the extremely acidic conditions of the symbiosome to exploit symbiont-produced sterols (Fig. 4G). We propose that whereas ubiquitously expressed canonical NPC2 homologues are ‘workhorses’ in sterol trafficking throughout the host, non-canonical NPC2s are spatiotemporally regulated to accumulate as the symbiosome matures, developing into a unique compartment optimized to promote the interaction and communication of symbiotic partners (Fig. 4G). This advantageous adaptation of existing molecular machinery allows symbiotic cnidarians to flexibly use symbiont-derived sterols, supporting survival in nutrient-poor environments. More broadly, our findings indicate that carbon acquisition by lipid transfer, similar to other symbioses [*Keymer*], is a major driver of coral-algal symbiotic relationships as means to adapt to various ecological niches by efficient exploitation of limited resources.

## Acknowledgements

We thank Britta Brügger, Christian Lüchtenborg, and Iris Leibrecht for assistance with sterol-binding lipidomics; Masayuki Hatta for *Acropora* coral and embryo collection and advice; Thomas Holstein for *Nematostella* provision, qPCR instrument access, and together with Sergio P. Acebrón for cell culture materials; Natascha Bechtoldt for qPCR technical assistance; Emmanuel Gaquerel, Anne Terhalle, and Michael Büttner for GC/MS instrument access and advice; Frauke Graeter, Agnieszka Obarska-Kosinska, Rebecca Wade, and Aura Navarro Quezada for input on theoretical comparative and evolutionary analyses. We thank Life Science Editors for editorial assistance with the manuscript. **Funding:** Support was provided to E.A.H., V.A.S.J., I.M., and A.G. by the Deutsche Forschungsgemeinschaft (DFG) (Emmy Noether Program Grant GU 1128/3-1), the European Commission Seventh Framework Marie-Curie Actions (FP7-PEOPLE-2013-CIG), the H2020 European Research Council (ERC Consolidator Grant 724715), the Boehringer Ingelheim Stiftung (Exploration Grant), and the future concept of Heidelberg University within the Excellence Initiative by German federal and state governments to A.G.; to V.A.S.J. by the European Molecular Biology Organization (Long-Term Fellowship); and to D.K. and T.S. by CellNetworks Heidelberg and the German Research Foundation (DFG, TRR83 and SFB1324) to Britta Brügger. **Author contributions:** E.A.H. and A.G. directed the research; E.A.H. primarily conducted the experiments and analyses with contributions from I.M. with immunoprecipitations, cloning, symbiont genotyping, and Western blots; V.A.S.J. conducted the immunofluorescence, confocal imaging, and IF quantification; D.K. designed and, together with E.A.H. and T.S., conducted and analyzed sterol-binding lipidomics. E.A.H. and A.G. wrote the manuscript. All authors approved the manuscript.

## Materials and Methods

### Computational Methods

#### NPC2 Bayesian consensus phylogeny construction

Genomes and, if available, proteomes and transcriptomes were downloaded from the sources in Supp. Table S1 and loaded into Geneious v.9 [*Kearse*]. If available, proteomes and transcriptomes were searched with BLASTp and BLASTx (both v.2.8.0), respectively, using as queries NPC2 homologues from *Aiptasia*, human, and relevant related and/or basal taxa. Each gene was located in its respective genome via discontinous Megablast, and the locus was manually annotated for complete gene (start and stop codon) and intron/exon structure. No NPC2 homologues were found in the single-celled eukaryotic filasterian *Capsaspora owczarzaki*; top BLASTp hits to NPC2 included two homologues of phospholipid transfer protein. With similar sizes to NPC2 and a predicted conserved ML superfamily domain, these were included in analyses and one (XP_004344261.1) used as an outgroup during phylogenetic tree construction as described below. In all sequences, signal peptide sequences were predicted using the SignalP4.0 server [*Petersen*] and, together with stop codons, removed from further analyses.

The genomes of 24 metazoans were included in the analysis, and phyla were represented by at least one species where available (when possible, marine taxa were chosen). Accession numbers and source information is given in Table S1. The resulting 77 NPC2 homologue sequences were imported into MEGA7 (v7.10.8) [*Kumar*] and aligned by codon using MUSCLE with default parameters and manually trimmed. The best model was calculated to be General Time Reversible with gamma distributed rate variation among sites (GTR+G). Bayesian phylogenies were inferred using MrBayes v.3.2.6 [*Huelsenbeck*] plugin in Geneious, with the GTR model, gamma rate variation, and five gamma categories. The consensus tree was estimated from four chains (temperature 0.2) for 1,000,000 generations, sampling every 200^th^ tree after 25% burn-in. The default prior was used: unconstrained branch lengths of GammaDir at 1, 0.1, 1, and 1.

#### Adaptive evolution

Evidence of diversifying and episodic selection on different sites among the NPC2 proteins were calculated using the DataMonkey server (http://classic.datamonkey.org) for the HyPhy program suite [*Kosakovsky Pond_1,2*]. Briefly, *Aiptasia* and *Nematostella* canonical and non-canonical NPC2 sequences were aligned by codon using MUSCLE in MEGA7 as above, and the best substitution model was calculated to be GTR+G+I for each. Bayesian phylogenies were inferred with MrBayes as above except for the following parameters: GTR+G+I, 4 gamma categories, 50,000 generations and sampling every 100^th^ tree after 20% burn-in. Trees were uploaded on the DataMonkey server and analysed with: i) fixed effects likelihood (FEL); ii) random effects likelihood (REL); iii) single-likelihood ancestor counting (SLAC) [*Kosakovsky_Pond 3*]; and iv) mixed effects model of evolution (MEME) [*Murrell*], and results of the tests were concatenated with the “Integrative Selection Analysis” tool. The program Rate4Site [*Mayrose*] was used to calculate per-site relative evolutionary rates. Briefly, protein sequences of *Aiptasia* and *Nematostella* canonical NPC2s or non-canonical NPC2s were aligned in MEGA7 using MUSCLE with default parameters, the termini trimmed, and the best substitution models calculated as WAG. With the alignments and user-generated trees as input, Rate4Site was run with the rates inferred using the empirical Bayesian method, a gamma prior distribution of 16 discrete categories, evolutionary model WAG, and branch lengths optimization with ML using a gamma model. The command was: *“*rate4site –s Alignment.fasta –t MrBayesTree.newick-o normalized_rates.txt −a Aiptasia1 (#or 6) –ib –mw −y original_rates.txt.”

### Live organism culture and collection

#### Aiptasia adults

Symbiotic and aposymbiotic *Aiptasia* adults were cultured as describes [*Grawunder*]; symbiotic animals rendered aposymbiotic [*Matthews*] were kept so for over one year before experimentation, with the exception of the “intermediate” aposymbiotic animal (Fig. 2). Animals were fed three times weekly with *Artemia* brine shrimp nauplii, shown to contain only cholesteroa [*Tolosa*]. Animals were then starved for at least four weeks prior to sampling. For sampling, animals were removed from their tanks simultaneously around mid-day, blotted briefly on lab tissue to remove excess seawater, and then prepared for either qPCR or GC/MS. For qPCR, animals were added to 1 ml Trizol (15596026, Life Technologies), after which they were quickly homogenized with a homogenizer (Miccra D-1, Miccra GmbH) at setting 3 for 10-15 seconds. Each tube was subsequently frozen at −80°C until RNA extraction. For GC/MS, animals were added to 400 µl ultrapure water, homogenized, and immediately processed.

#### Aiptasia eggs and larvae

Adults of strains F003 and CC7 were cultured and induced to spawn as described [*Grawunder*]. For GC/MS, approx. 1000-3000 unfertilized eggs from female-only tanks were collected gently with transfer pipette within 2 h of spawning, washed once quickly in water and once quickly in methanol, and resuspended in 750µl methanol. For NPC2 immunofluorescence (IF) during symbiosis establishment, *Aiptasia* larvae 2 days post-fertilization (dpf) at a density of 300-500/ml FASW were exposed to *Symbiodinium* strain SSB01 as described [*Bucher*] at 10,000 cells/ml. Larvae and algae were co-cultivated for 3 d, until at 5 dpf the larvae were filtered, washed, and resuspended in fresh FASW at a density of 300-500/ml. Larvae were fixed at the indicated time-points with 4% formaldehyde in filtered artificial seawater (FASW) rotating for 45 min RT, washed twice with PBT (1x PBS pH 7.4 + 0.2% Triton-X), and stored in PBS at 4°C in the dark.

#### Nematostella adults

For qPCR, stock cultures of mixed-sex *Nematostella* were kept in 12:12 L:D at 26°C and fed weekly with *Artemia* nauplii. Animals were then separated and either starved or fed *Artemia* nauplii daily for 14 d with subsequent daily water changes; animals were then starved for a further 2 d and then sampled as for *Aiptasia* qPCR. For GC/MS, mixed-sex *Nematostella* were kept in constant dark at 16°C, fed once weekly with *Artemia* nauplii, and water changed the following day; animals were starved for 10 d and then sampled as for *Aiptasia* GC/MS.

#### Acropora digitifera adults, larvae, and primary polyps

Colonies of the coral *Acropora digitifera* were collected off Sesoko Island (26°37’41”N, 127°51’38”E, Okinawa, Japan) according to Okinawa Prefecture permits and CITES export and import permits (T-WA-17-000765). Corals were kept as described [*Wolfowicz*] in tanks with running natural seawater and under partially shaded natural light at Sesoko Tropical Biosphere Research Center (University of Ryukyus, Okinawa, Japan). Colonies were isolated prior to spawning, and subsequently-spawned bundles of symbiont-free gametes were mixed for fertilization of defined crosses. The resulting planula larvae were maintained at approximately 1000 larvae/L in 10μm-filtered natural seawater (FNSW) exchanged daily. For GC/MS, samples from spawning on the evening of 27 May 2016 were collected from adult parental colonies and their embryo offspring 19 and 24 hpf, respectively, and immediately transferred to methanol. For qPCR, adults and their embryos from spawning on the evening of 6 June 2017 were simultaneously collected at the indicated hpf and immediately transferred into RNAlater (AM7020, Thermo Fisher Scientific). All samples were transferred to 4°C within several hours and then to −20°C within 2 d, where they were kept until processing. To generate juvenile primary polyps, larvae were induced to settle at 6 dpf and infected with *Symbiodinium* strain SSB01 as described [*Wolfowicz*] for 4 d. Resident *Symbiodinium* in adult parental colonies were typed as previously described [*Grawunder*]: 10 bacterial clones were sequenced per coral colony and all were identical, identified by BLASTn to the nr NCBI database as *Symbiodinium* Clade C1.

#### Symbiodinium cultures

Clonal and axenic *Symbiodinium* of the strains SSB01 (clade B), SSA01 (clade A), SSA02 (clade A), and SSE01 (clade E) [*Xiang*] as well as the non-clonal, non-axenic strains CCMP2466 (clade C) and CCMP2556 (clade D) purchased from the National Center for Marine Algae and Microbiota (NCMA, Bigelow Laboratory for Ocean Sciences, Maine, USA) were cultured as described [*Wolfowicz*]. For GC/MS, 2.6×10^7^ cells were collected at mid-day by centrifugation of cultures at 1000xg at RT, washed twice in FASW, and the cell pellet resuspended in ultrapure water and processed as described. Because of its use in GC/MS-based sterol profile comparisons to the endogenous *A. digitifera* symbiont, strain CCMP2466 (clade C) was re-typed as previously described [*Grawunder*]; from a single amplification reaction, 2 bacterial clones were sequenced and had best BLAST hit in NCBI nr database to *Symbiodinium* in Clade C.

#### Cell culture

HEK 293T cells were cultured in 1X DMEM medium (41966029, Gibco/Thermo Scientific) with 10% FBS and 1% pen/strep (100µg/ml final concentrations). Cells were grown at 37°C with 5% carbon dioxide and passaged regularly.

### Gene expression

#### RNA extraction and qPCR

RNA was extracted according to a hybrid protocol [*Polato*] with phenol-chloroform and the RNeasy kit (74104, Qiagen). RNA was qualitatively and quantitatively assessed via gel electrophoresis and NanoDrop spectrophotometry (Nanodrop1000), respectively, aliquoted and flash frozen in liquid nitrogen and stored at −80C. First strand cDNA synthesis was performed with the ReadyScript cDNA synthesis kit (RDRT, Sigma Aldrich) according to the manufacturer’s instructions. qPCR was performed in 96 well plate format, with each reaction containing 0.4 µm each primer, 50 ng cDNA, and 1X SensiFast SYBR Hi-ROX qPCR master mix (BIO-92005, Bioline) in 20 µl total; reactions were measured on a StepOnePlus (Applied Biosystems). The gene encoding 40S Ribosomal Protein S7 (RPS7) was chosen as a comparison/baseline gene due to its demonstrated stability in a previous study [*Lehnert*]. Primers are in Supp. Table S2; all primer pairs were validated by amplicon sequencing through either TOPO-TA cloning (450071, Thermo Fisher Scientific) according to the standard protocol or, for *Acropora* and *Nematostella*, direct sequencing of qPCR products, with at least three sequences per product. Additionally, melt curves performed after each qPCR run confirmed the existence of single products per reaction. Amplification efficiencies of each primer pair were determined by a 3-or 4-point dilution series. Output was analyzed with the Bayesian analysis pipeline MCMC.qPCR [*Matz*] run according to standard protocol (https://matzlab.weebly.com/data--code.html). For *Acropora* adults and embryos, the model was run with ‘naïve’ parameters. For comparative expression within symbiotic *Aiptasia* and fed *Nematostella*, the analysis was run with ‘informed’ parameters setting RPS7 as a reference gene. Log_2_ (fold change) and Log_2_ (transcript abundance) were determined from command ‘HPDsummary’ with and without ‘relative=TRUE’, respectively; p-values of differential expression were calculated with command ‘geneWise’ on the former.

#### Nematostella embryonic development

Expression data on *Nematostella* embryonic development and comparative adults were obtained from NvERTx [*Warner*] (http://ircan.unice.fr/ER/ER_plotter/home). Transcripts were identified by BLAST search to the NvERTx database as the *NPC2* homologues XM_001622824.1 (NvERTx.4.51280); XM_001627305.1 (NvERTx.4.192779); XM_001635452.1 (NvERTx.4.142169), and the *RPS7* homologue jgi|Nemve1|54581|gw.133.186.1 (NvERTx.4.145315). Transcript abundance counts at 0 h post-fertilization (unfertilized) were taken from the Counts table, comprising duplicate samples of 300 embryos per sample [*Fischer*]. As a baseline for typical gene expression in adults, transcript abundance counts were taken from the “uncut controls” (UC) in an adult regeneration experiment, comprising triplicate samples of untreated 6-week-old adults, 300 animals per sample [*Warner*].

### Sterol profiling with gas chromatography/mass spectrometry (GC/MS)

Samples were extracted with a modified Bligh-Dyer method: briefly, either 300 µl aqueous animal homogenate (*Aiptasia* or *Nematostella* adults) was added to 750 µl HPLC-grade methanol, or 300 µl ulta-pure water was added to the sample already in 750 µl methanol or ethanol (*Acropora*). The mixture was incubated shaking at 70°C for 45 min, combined with 375 µl HPLC-grade chloroform and 300 µl ultra-pure water, thoroughly mixed, and centrifuged for 10 min at 4°C at 6000xg. The lower organic phase was then dried fully by vacuum centrifugation at ≤ 45°C and then saponified by adding 500 µl of 5% KOH in a 9:1 methanol:water solution and incubating at 68°C for 1 h. The mixture was then combined with 250 µl water and 750 µl chloroform, centrifuged 10 min at 6000xg, and the organic phase was isolated and dried to completion with a SpeedVac. Lipids were derivatized to trimethylsilyl ethers with 25-40 µl MSTFA in Pyridine (#69479, Sigma Aldrich) at 60°C for 0.5-1 h and immediately analyzed. 1 µl of each mixture was injected into a QP2010-Plus GC/MS (Shimadzu) and with a protocol (adapted from [*Schouten*]) as follows: oven temperature 60°C, increase to 130°C at 20°C/min, then increase to 300°C at 4°C/min and hold for 10 min. Spectra were collected between m/z 40 and 850 and data were analyzed in GCMS PostRun Analysis Software (Shimadzu). Spectra were compared to the National Institute of Standards and Technology 2011 database for matches; retention time relative to cholesterol and high-confidence database matches, were used to assign sterol identities and matching sterols between samples. Relative sterol composition as percent of total sterols were calculated from integrated peak intensity on the total ion chromatograph for each sample.

### *Aiptasia*-specific anti-NPC2 antibodies

Antibodies were raised against the peptides K-YGIDVFCDEIRIHLT (XM_021052412), K-AKNDIFCNSIPFNLV (XM_021052404), and K-VQNNVLCGEVTLTLM (XM_021052381) coupled to the adjuvant keyhole limpet hemocyanin in rabbits (BioScience GmbH). Antibodies were affinity-purified from the antisera with the peptides coupled to NHS-Activated Sepharose Fast Flow 4 (17090601, GE Health Care Life Sciences) according to the manufacturer’s protocols.

### Western blots of *Aiptasia* and *Symbiodinium* homogenates

Two aposymbiotic or symbiotic adult *Aiptasia* were homogenized in buffer A with 2X Halt Protease Inhibitor Cocktail (78430, Thermo Fisher Scientific) and then sonicated on ice (Sonifier 250, Branson Ultrasonics) with two rounds of 25 pulses at duty cycle 40%, output control 1.8. From a culture of *Symbiodinium* strain SSB01, 1.2×10^7^ cells were collected by centrifugation at 6,000xg at RT. After addition of buffer A and glass beads (425-600 µm), cells were disrupted by vortexing six times for 1 min each, with 1 min on ice in between each. Homogenates were then transferred to a new tube with syringe (G23 needle) for further disruption. All homogenates were then centrifuged at 20,000*xg* for 10 min at 4°C, and three sets of identical volumes of the supernatants were resolved on a 12% Tricine-SDS-Page gel and transferred by Western blot onto nitrocellulose membranes. Membranes were blocked for 1 h in 5% milk PBS-T and then incubated with antibodies raised against three different non-canonical *Aiptasia* NPC2s (XM_021052404 at 1:4000, XM_021052412 at 1:1000 and XM_021052381 at 1:500) and anti-tubulin (T90026, Sigma-Aldrich) as loading control at 1:1000 in 5% milk PBS-T at 4°C overnight, followed by incubation with HRP-coupled anti-rabbit and HRP-coupled anti-mouse (Jackson ImmunoResearch) at 1:10000 in 5% milk PBS-T at RT for 1 h, and then detection with ECL (GERPN2232, Sigma-Aldrich) and imaging on ECL Imager (ChemoCam, Intas).

### Immunoprecipitation-lipidomics of NPC2-sterol binding

#### Cell culture lysates and symbiont extracts

For recombinant expression, either a canonical (XM_021041171) or a non-canonical (XM_021052404) *Aiptasia* NPC2 protein with the signal peptide and stop codon removed were cloned behind the cytomegalovirus promoter in a pCEP-based vector followed by a five-proline linker, cleavage-resistant mCherry (crmCherry) [*Huang*], and 3xHA tag (YPYDVPDYA); a control vector contained only crmCherry:3xHA. Vectors were transiently transfected with Lipofectamine2000 (11668019, Invitrogen/Thermo Fisher Scientific) according to the manufacturer’s protocol into HEK cells in 10 cm diameter dishes. Cells were harvested after growth for 48 h at 32°C by rinsing with PBS, addition of 1 ml of buffer A, B, C, or D (see Buffers below) with Halt Protease Inhibitor Cocktail at 2X (78430, Thermo Fisher Scientific), and scraping with a cell scraper. The resulting lysates were then sonicated on ice (Sonifier 250, Branson Ultrasonics) with two rounds of 25 pulses at duty cycle 40%, output control 1.8, centrifuged at 20,000xg for 20 min at 4°C, and supernatants used in binding assays. Approximately 2.5×10^8^ cells of *Symbiodinium* strain SSB01 [*Xiang*] approx. 7 d after passaging were collected by centrifugation at 6000xg at 22°C. Cells were washed in 10 ml of Buffer A, B, C, or D (per the corresponding HEK cell lysate), and then 5ml buffer was added to the pellet and cells were sonicated twice for 5 min at duty cycle 80%, output control 3. We noted that extracts were best when allowed to heat slightly (not boiling) during sonication. Extracts were centrifuged at 6000xg for 10 min at RT, and supernatants used in binding assays.

#### Immunoprecipitation

Cell lysates were incubated together with symbiont extracts (450 µl and 500 µl, respectively) for 30 min at room temperature (21-23°C) rotating, after which 25 µl anti-HA beads (130-091-122, Miltenyi Biotech) were added and the mixtures incubated rotating at RT for a further 30 min. Mixtures were then passed through magnetic columns on a magnetic plate (130-042-701, Miltenyi Biotech) pre-rinsed with 200 µl of the corresponding Buffer A, B, C, or D. Bound material on the column was rinsed four times with Buffer A, B, C, or D with protease inhibitor, and then once with the corresponding buffer half-diluted and without protease inhibitor. Lipids were eluted by application of 20 µl HPLC-grade methanol to the column for 5 min incubation, followed by 100 µl methanol and collected into HPLC glass bottles with glass inserts and Rubber/PTFE caps (Neochrom, NeoLab). Eluates were immediately transferred to ice and then −20°C until lipidomics processing on the same day. Proteins were then eluted by application of 20 µl loading dye (20 mM DTT, 60 mM Tris pH 6.8, 20% glycerol, 1% SDS, 0.01% Bromophenol blue) at 100°C to the column for 5 min incubation, followed by 50 µl loading dye and collection. Samples were then heated to 95°C for 3 min and then immediately resolved by SDS PAGE on 10% Tris-Glycine gels and transferred by Western blot onto nitrocellulose membranes. Membranes were blocked for 1 h in 5% milk TBS-T and then incubated with anti-mCherry (PA5-34974, Thermo Fisher Scientific) at 1:3000 in 5% milk TBS-T at 4°C overnight, followed by incubation with conformation-specific HRP-coupled anti-rabbit (5127S, Cell Signaling Technology) at 1:2000 in 5% milk TBS-T at RT for 2 h, and then detection with ECL (GERPN2232, Sigma-Aldrich) and imaging on ECL Imager (ChemoCam, Intas).

#### Immunoprecipitation from Aiptasia homogenates

Purified polyclonal antibody against XM_021052404 (described above) was coupled to epoxy magnetic beads in the Dynabeads Antibody Coupling Kit (14311, Thermo Fisher Scientific) per the manufacturer’s instructions. Beads (1 mg per reaction) were then incubated with homogenates of *Aiptasia* CC7 for 16 h rotating at 4°C; control reactions contained uncoupled beads. After washing, protein-lipid complexes were immunoprecipitated via magnetic separation and eluted from beads with 200 mM glycine, pH 2.3, then neutralized with 0.1 M Tris-HCl, pH 8.5. An aliquot was taken for Western blot visualization of proteins; the remainder was extracted for lipids with a mixture of chloroform:methanol:water (final ratios 8:4:3). After 5 min centrifugation at 3000x*g*, the organic phase was isolated and dried completely by vacuum centrifugation at 30°C for 1.5 h, and then reconstituted with 100 µl methanol and collected into HPLC glass bottles with glass inserts and Rubber/PTFE caps (Neochrom, NeoLab). Eluates were immediately transferred to ice and then −20°C until lipidomics processing on the same day.

#### Buffer

**A:** 200 mM Ammonium Acetate, pH 7; **B**: 200 mM Ammonium Acetate, pH 5; **C**: 50 mM MES, 150 mM NaCl, 0.004% Nonidet P-40; **D**: 50 mM Tris, 150 mM NaCl, 0.004% Nonidet P-40, pH 7.5.

#### Lipidomics

50 µl of each eluate was added to chloroform-rinsed glass tubes, followed by addition of 100 pmol cholesterol-D6 as an internal standard. Samples were dried under nitrogen, derivatized with addition of 50 µmol acetylchloride in methylene (708496, Sigma Aldrich) for 30 min at RT, and then dried under nitrogen again. Samples were then dissolved in 100 mM ammonium acetate in methanol and loaded into 96-well plates for analysis. A standard curve in duplicate of pmol cholesterol/stigmasterol at 50/25, 250/125, 500/250 was always processed in parallel. Samples were injected by a TriVersa NanoMate autosampler (Advion) held at 10°C on nanospray mode with positive polarity at 1.2 kV and 0.4 psi gas pressure through a D-Type silicon chip with 4 µm nominal diameter. Samples were analysed on a QTRAP 5500 (SCIEX) Hybrid Triple-Quadrupole/Linear Ion Trap at 60°C in positive ion and neutral loss scan mode (loss of acetate, 77.05 Da), with low Q1 resolution and high Q3 resolution at a scan rate of 200 Da/sec and 120 scans total. The DMS (differential mobility spectrometer) to select for ion mobility was enabled and ran at 60°C, medium pressure (24 psi), and a compensation voltage (COV) of 4.4 kV. In every run a pooled mixture of all samples was run with a COV ramp from 0 to 20 kV to confirm the appropriate COV. The machine was driven by Analyst software version 1.6.3 (SCIEX), and output was processed in LipidView software to detect and quantify sterols by peak intensities. Sterol concentrations were calculated by normalization to the cholesterol-D6 internal standard, subtraction of blank samples, and comparison to the standard curve.

### Immunofluorescence of NPC2 in *Aiptasia* larvae

Fixed larvae in PBS at 4°C were permeabilized in PBT for 2 h at RT. Samples were then incubated in blocking buffer (5% normal goat serum and 1% BSA in PBT) overnight at 4°C and then with primary antibody diluted in block buffer for 4 h at RT at the following concentrations: 45 μg/ml (XM_021052404), 15 μg/ml (XM_021052412), and 20 μg/ml (XM_021052381). Samples were then washed twice for 5 min with PBT at RT, twice for approx. 18 h at 4°C, then incubated with secondary antibody (goat anti-rabbit IgG-Alexa488; ab150089, Abcam) diluted to 4 μg/ml in block buffer for approx. 5 h at RT. Samples were then washed with PBT three times for 5 min each at RT, then approx. 18 h at 4°C. Sample were then incubated with Hoechst 33342 at 10 μg/ml in PBT for 1 h at RT, washed 3x with PBT for 5 min each, and then washed into PBS at 4°C overnight. PBS was replaced with 95% glycerol with 2.5 mg/ml DABCO, and the larvae were mounted for microscopy.

### U18666A exposure in *Aiptasia* and *A. digitifera*

Adult *Aiptasia* polyps were added to 6-well culture plates and allowed to attach for 2 d before exposure to U18666A (U3633, Sigma Aldrich) in DMSO at the indicated concentrations in FASW, where the final percentage of DMSO in the FNSW was <0.05%. Polyps were cultured at 26°C at 12:12 L:D and daily photographed followed by wash and drug re-addition. *Acropora* polyps hosting *Symbiodinium* SSB01 were exposed to U18666A as described, except that they were cultured in FNSW.

### Microscopy

Confocal microscopy of NPC2 immunofluorescence was performed using a Leica SP8 system with and HC PL Apo CS2 63x/1.30 GLYC objective. Hoechst was excited at 405 nm and detected at 410-501 nm, and algal autofluorescence was excited at 633 nm and detected at 645-741 nm. In a second sequential scan, Alexa-488 (secondary antibody) was excited at 496 nm and detected at 501-541 nm. Z-stacks were collected with a step size of 0.5 μm and 3x line averaging. A zoom factor of 5 or, for whole larvae, 1.33, was used, and a pinhole of 1 Airy unit. Quantification and imaging NPC2 IF over a time-course was carried out using a Nikon Eclipse Ti epifluorescence compound microscope with a Plan Apo λ 40x objective, Sola light source, and GFP filter set. Images were captured with a Nikon DS-Qi2 with an exposure time of 1 s. Three replicates at each time-point were quantified, with quantification proceeding until 50 infected larvae per replicate were counted. At least three replicates per time-point were carried out. Fluorescence microscopy of *Aiptasia* adults was carried out using a Nikon SMZ18 fluorescence stereoscope with a 0.5X objective; endogenous autofluorescence of *Symbiodinium* photosynthetic antennae was visualized with a Texas Red filter set, and images were captured at magnification 15X with an Orca-Flash4.0 camera (C11440, Hamamatsu) at 300 ms exposure using Nikon Elements software. *Acropora* polyps were photographed as described [*Wolfowicz*].

## Supplementary Material

**Supp Fig. S1.**
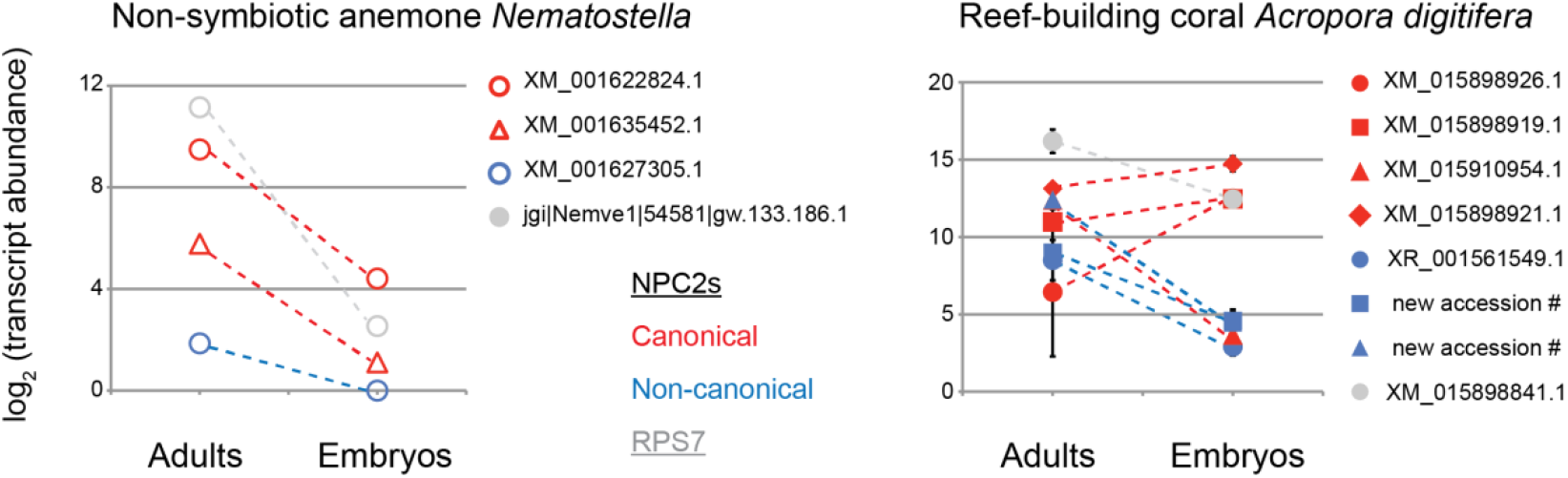
Differential maternal loading of canonical and non-canonical NPC2 transcripts in embryos of *Acropora* and *Nematostella*. **Left,** Expression in *Nematostella vectensis* adults and in embryos. From RNAseq regeneration/development dataset on NvERTx server [*Warner, Fischer, Helm*]. **Right,** Expression in both *Acropora digitifera* parent colonies and their offspring immediately after spawning. qPCR average of two biological replicate samples, each sample in technical duplicate. Average value ±SD (error bars). Difference between non-canonical NPC2s in adults and embryos significant, Student’s paired *t*-test, p=0.007 (canonical, p=0.18).

**Supp Fig. S2.**
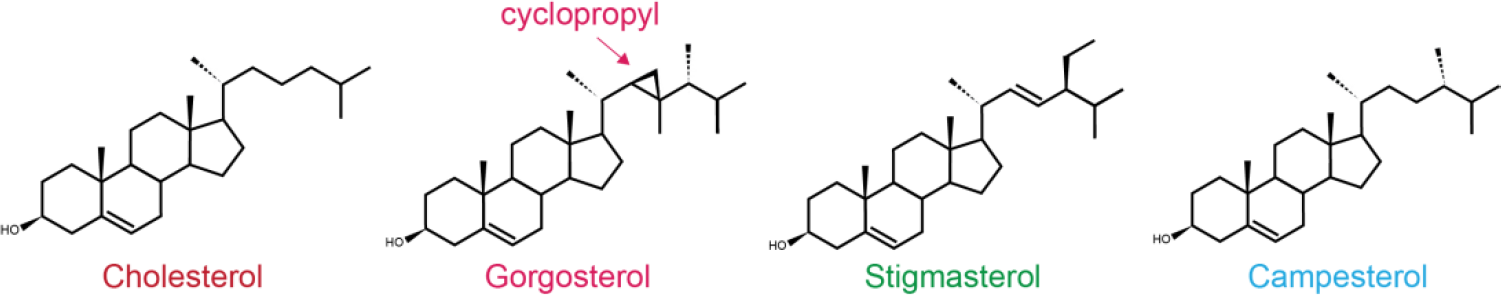
Some of the symbiont-produced sterols. Symbionts, like their dinoflagellate relatives, produce many sterols, including the unique gorgosterol with its unusual cyclopropyl group. Colors correspond to those in Fig. 3 A, C.

**Supp Fig. S3.**
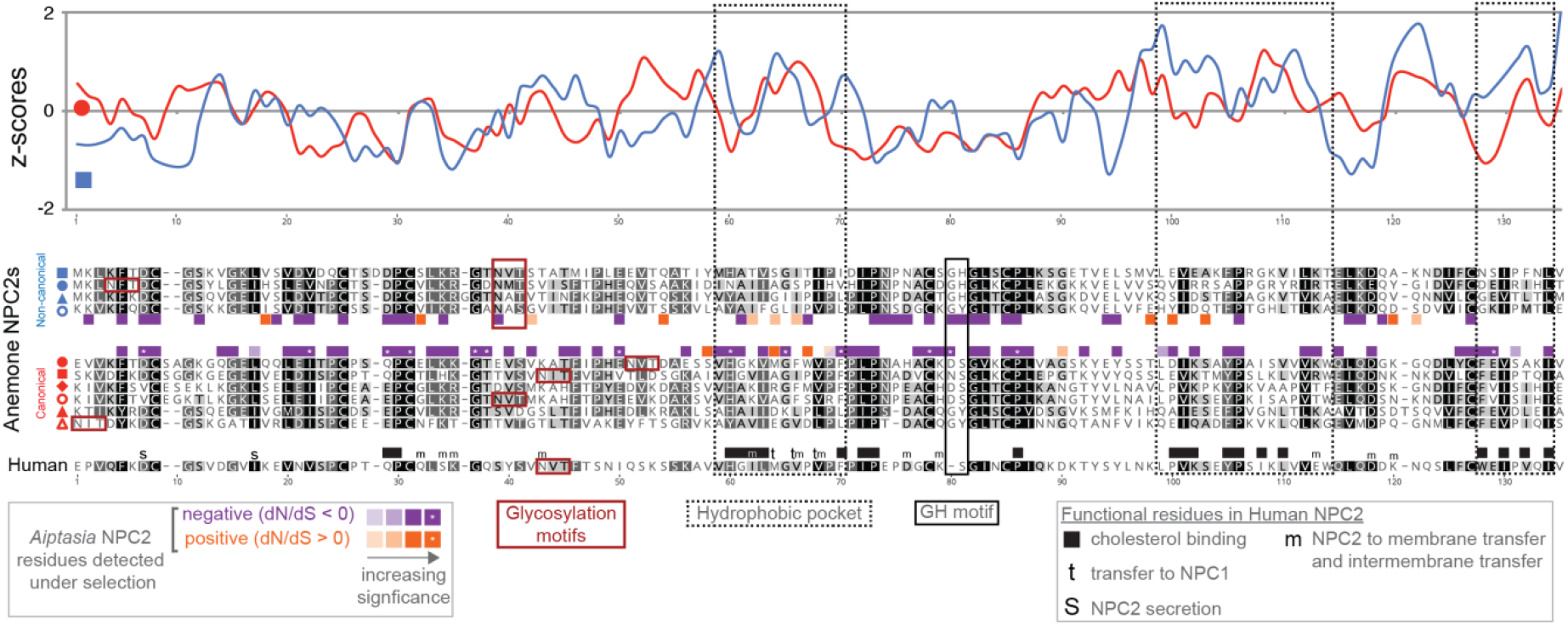
Additional adaptive evolution analysis of *Aiptasia* canonical and non-canonical NPC2s. Relative evolutionary rates per residue for the investigated *Aiptasia* canonical (XM_021041171) and non-canonical (XM_021052404) proteins mapped onto the sequence alignments from Fig. 3A. Relative rates were calculated with Rate4Site [*Mayrose*] using sequence alignments of *Aiptasia* and *Nematostella* canonical or non-canonical proteins (see Methods). Conservation scores from both outputs were transformed to z-scores, which were smoothened by averaging within a sliding-scale window of 3 residues.

**Supp Fig. S4.**
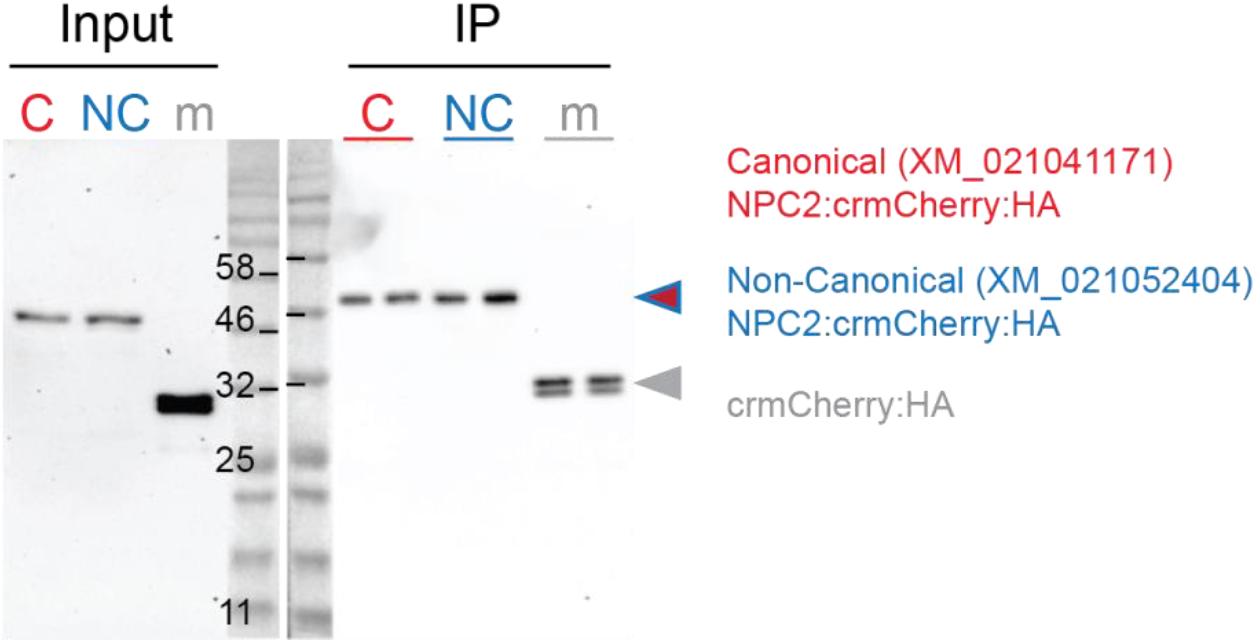
Immunoprecipitation-lipidomics: protein inputs and immunoprecipitation (IP) elutions. HEK cell lysate input containing recombinant NPC2s or crmCherry control, and subsequent protein elutions of anti-HA IP reactions, in duplicate. After washing, lipids were eluted (Fig. 3C-D) from IP reactions prior to protein elution shown here.

**Supp Fig. S5.**
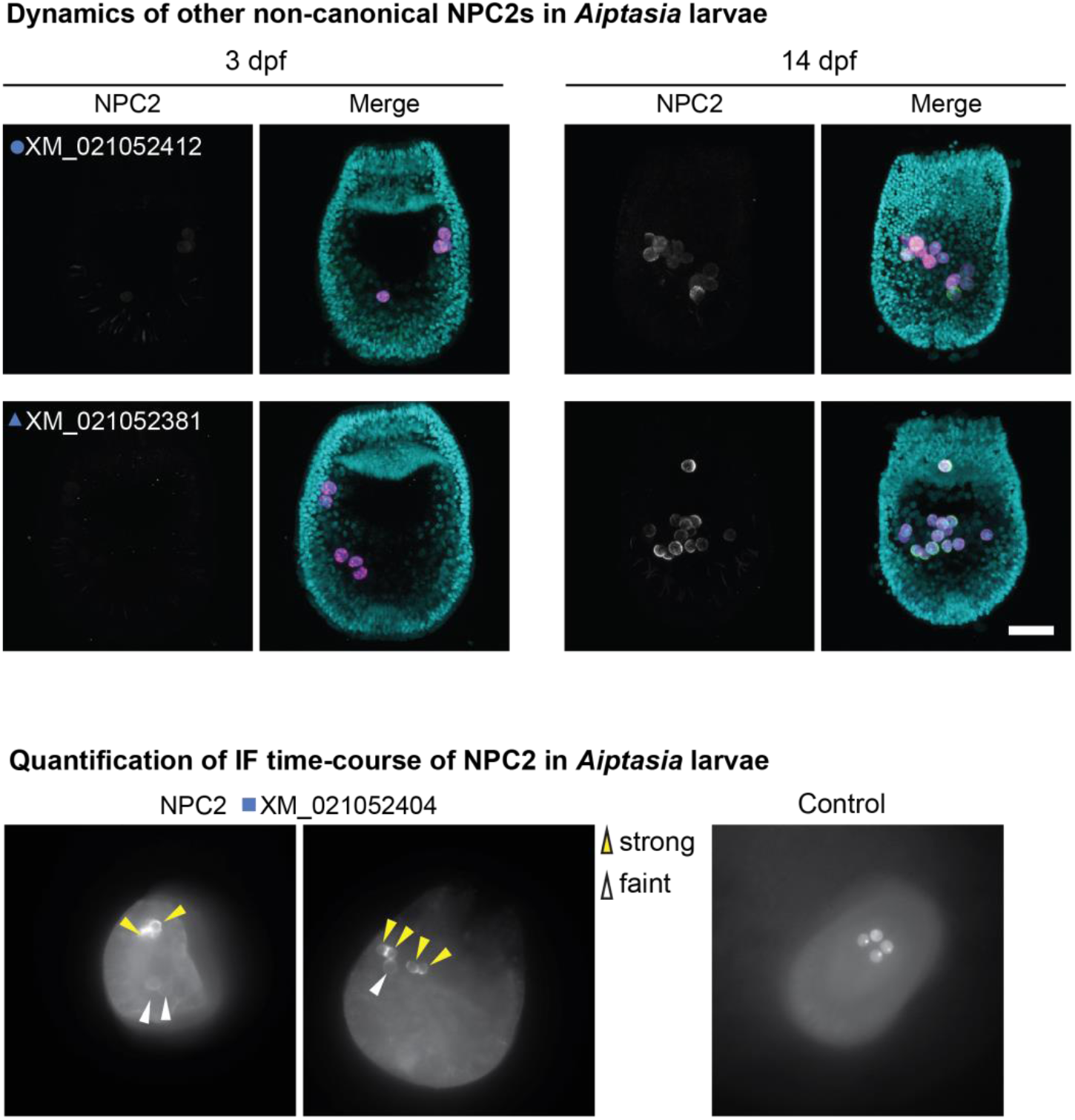
Dynamic recruitment of other non-canonical NPC2s to intracellular symbionts increases as symbiosis matures. **Top,** Immunofluorescence (IF) of non-canonical NPC2s in *Aiptasia* larvae containing intracellular symbionts of *Symbiodinium* strain SSB01 at 3 and 14 dpf. Merge channels as in Fig. 4. Control, secondary antibody only. Scale bar, 25 μm. **Bottom,** Example images for quantification of NPC2 IF time-course in Fig. 4C-D, with strong and weak NPC2 staining indicated by arrows. Control, secondary antibody only.

**Supplementary Table S1.**

| Species | Genome Version | Genome Accession Number (Source, if not NCBI) | Proteome (P), Transcriptome (T), Gene Models (GM), if different from genome (Source, if not NCBI) |
| --- | --- | --- | --- |
| *Acropora digitifera* | 1.1 | GCA_000222465.2_Adig_1.1_ | T: GCF_000222465.1_Adig_1.1_rna.fna |
| *Adineta vaga* | 1 | GCA_000513175.1_AMS_PRJEB1171_v1 | P: Adineta_vaga.v2.pep.fa |
| *Aiptasia (Exaiptasia) pallida* | 1.1 | GCF_001417965.1_Aiptasia_genome_1.1 |  |
| *Amphimedon queenslandica* | 1.0 | GCF_000090795.1_v1.0 |  |
| *Aplysia californica* | 3.0 | GCF_000002075.1_AplCal3.0 |  |
| *Branchiostoma floridae* | 2 | GCF_000003815.1_Version_2 |  |
| *Capitella teleta* | Capcal | GCA_000328365.1_Capcal |  |
| *Capsaspora owczarzaki* | 2 | GCA_000151315.2_C_owczarzaki_V2 |  |
| *Ciona intestinalis* | KH | GCF_000224145.2_KH |  |
| *Crassostrea gigas* | 9 | GCF_000297895.1_oyster_v9 |  |
| *Daphnia magna* | 2.4 | GCA_001632505.1_daphmag2.4 |  |
| *Homo sapiens* | GRCh38.p12 | GCA_000001405.27 | Genome Data Viewer<br>(https://www.ncbi.nlm.nih.gov/genome/gdv/) |
| *Hydra vulgaris* | 2.0 | Hm105_Dovetail_Assembly_1.0.fa<br>(https://research.nhgri.nih.gov/hydra/) | GM: hydra2.0_genemodels<br>T: Juliano_transcript; Petersen_transcript<br>(https://research.nhgri.nih.gov/hydra/) |
| *Hydractinia echinata* | draft | Run03Super.fa.gz<br>(https://research.nhgri.nih.gov/hydractinia/) | described in Török et al. 2016 |
| *Limulus polyphemus* | 2.1.2 | GCF_000517525.1_Limulus_polyphemus-2.1.2 |  |
| *Lingula anatine* | 1.0 | GCF_001039355.1_LinAna1.0 |  |
| *Macrostomum lignano* | 2 | GCA_001188465.1_ML2 | MLRNA150904.fa (www.macgenome.org) |
| *Montastrea cavernosa* | version March 2018 | Mcavernosa_03012018.fasta<br>(Yi Liao & Mikhail Matz, Univ. Texas-Austin, Zachary Fuller & Molly Przeworski, Columbia Univ. https://matzlab.weebly.com/data--code.html) | T: Mcavernosa.maker.transcripts.fasta<br>P: Mcavernosa.maker.proteins.fasta<br>(https://matzlab.weebly.com/data--code.html) |
| *Mnemiopsis leidyi* | MneLei_Aug2011 | GCA_000226015.1_MneLei_Aug2011 | P: unfiltered MLRB2.2.aa<br>GM: ML2.2.nt; T: ML_Cufflinks_transcripts.fa, ML_Trinity_transcripts.fa<br>(http://research.nhgri.nih.gov/mnemiopsis/) |
| *Nematostella vectensis* | 1 | GCA_000209225.1_ASM20922v1 |  |
| *Priapulius caudatus* | 5.0.1 | GCF_000485595.1_Priapulius_caudatus-5.0.1 |  |
| *Saccoglossus kowalevskii* | 1.1 | GCA_000003605.2_Skow_1.1 |  |
| *Strongylocentrotus purpuratus* | 4.2 | GCF_000002235.4_Spur_4.2 |  |
| *Trichoplax adhaerens* | 1.0 | GCF_000150275.1_v1.0 |  |

**Table S2.**
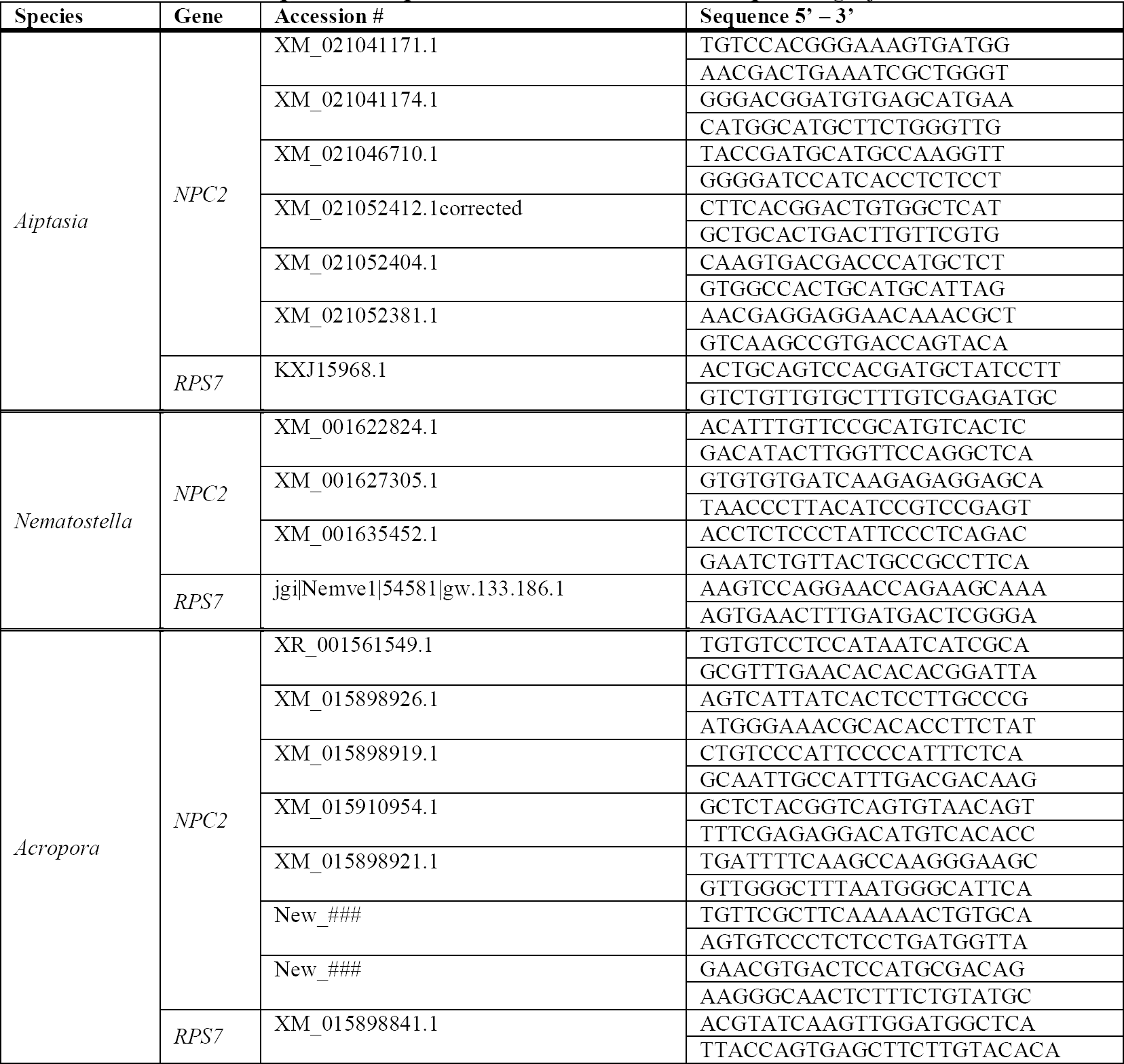
Primers for qPCR in *Aiptasia*, *Nematostella*, and *Acropora digitifera*

